# The Genealogical Sorting Index and species delimitations

**DOI:** 10.1101/036525

**Authors:** David J. Winter, Steven A. Trewick, Jon M. Waters, Hamish G. Spencer

## Abstract

The Genealogical Sorting Index (*gsi*) has been widely used in species-delimitation studies, where it is usually interpreted as a measure of the degree to which each of several predefined groups of specimens display a pattern of divergent evolution in a phylogenetic tree. Here we show that the *gsi* value obtained for a given group is highly dependent on the structure of the tree outside of the group of interest. By calculating the *gsi* from simulated datasets we demonstrate this dependence undermines some of desirable properties of the statistic. We also review the use of the *gsi* delimitation studies, and show that the *gsi* has typically been used under scenarios in which it is expected to produce large and statistically significant results for samples that are not divergent from all other populations and thus should not be considered species. Our proposed solution to this problem performs better than the *gsi* in under these conditions. Nevertheless, we show that our modified approach can produce positive results for populations that are connected by substantial levels of gene flow, and are thus unlikely to represent distinct species. We stress that the properties of *gsi* made clear in this manuscript must be taken into account if the statistic is used in species-delimitation studies. More generally, we argue that the results of genetic species-delimitation methods need to be interpreted in the light the biological and ecological setting of a study, and not treated as the final test applied to hypotheses generated by other data.

## Introduction

Genetic sequence data and phylogenetic methods are increasingly being used to aid in the discovery and delimitation of species (reviewed in Fujita *et al*. 2012). The widespread application of such data and analyses to alpha taxonomy has confirmed that evolutionarily distinct species will not necessarily fall into reciprocally monophyletic groups in phylogenies estimated from DNA sequences. Indeed, species can remain paraphyletic with respect to their close relatives in gene trees even millions of years after they begin to diverge (Tajima 1983; Hudson & Coyne 2002).

A number of methods have been developed with the objective of delimiting species using such unsorted gene trees (Knowles & Carstens 2007; O’Meara 2010; Yang & Rannala 2010, 2014; Ence & Carstens 2011; Zhang *et al*. 2011). The increasingly popular use of these methods in empirical species-delimitation studies has inspired a number of methodological papers exploring their statistical properties. These theoretical investigations have shown the methods to be powerful and accurate when their underlying assumptions are met, but it has become clear that violations of these assumptions can generate misleading results (Reid & Carstens 2012; Carstens *et al*. 2013; Edwards & Knowles 2014; Olave *et al*. 2014). Thus, species delimitation methods are most useful when their statistical properties are understood, and studies can be designed and interpreted in the light of these properties.

Although not exclusively designed for species-delimitation studies, the *gsi* of Cummings *et al*. (2008) has been widely used in this context (see references in Table 1). This statistic is a measure of the degree to which a pre-defined group of leaves in a phylogenetic tree falls into an exclusive region in that tree. The value of the statistic ranges from 0 to 1, with higher values corresponding to more phylogenetic exclusivity. In this way, the *gsi* aims to bridge the gap between monophyly and paraphyly as categorical terms, and in so doing, quantify the degree to which a lineage has become distinct as a result of evolutionary divergence. The calculation of the *gsi* is usually accompanied by an hypothesis test, in which the *gsi* for each group is compared to values calculated from trees in which the tip labels have been permuted.

**Table 1.** Summary of papers in which the gsi is principally used for species delimitation. “n.groups” refers to the greatest number of putative species considered in a single analysis

| Reference | Evidence for group assignment | n.<br>groups |
| --- | --- | --- |
| Martinsson *et al.* In press | Clades in mitochondrial gene tree | 2 |
| Sánchez-Ramírez *et al.* In Press | Existing taxonomic distinction, clades in gene trees | 17 |
| Gehesquière *et al.* In Press | Clustering of haplotypes | 2 |
| Egea *et al.* 2016 | Mitochondrial clades | 5 |
| Bon *et al.* 2015 | Existing taxonomic distinction | 3 |
| Hu *et al.* 2015 | Clades identified in other trees | 7 |
| Esposito *et al.* 2015 | Clustering of haplotypes | 11 |
| Su *et al.* 2015 | Clustering of haplotypes | 3 |
| Bagley *et al.* 2015 | Clades, other delimitation methods, existing taxonomic distinction | 10 |
| Vigalondo *et al.* 2015 | Clades in multilocus phylogeny and existing taxonomic distinction | 4 |
| Pažoutová *et al.* 2015 | Existing taxonomic distinction | 4 |
| Ramírez *et al.* 2014 | Existing taxonomic distinction | 2 |
| Fourie *et al.* 2014 | Existing taxonomic distinction and geographically isolated populations | 17 |
| Fernández-Mazuecos & Vargas 2014 | Existing taxonomic distinction | 6 |
| Viricel & Rosel 2014 | Morphology, clustering of genotypic data and geographic distribution | 2 |
| Derkarabetian & Hedin 2014 | Morphology, mitochondrial clades | 11 |
| Wang *et al.* 2014 | Existing taxonomic distinction, clades in gene trees | 3 |
| Ashalakshmi *et al.* 2014 | Existing taxonomic distinction | 4 |
| Udayanga *et al.* 2014 | Existing taxonomic distinction and congruent clades among gene trees | 10 |
| Almendra *et al.* 2014 | Existing taxonomic distinction and clades in gene trees | 3 |
| Lu *et al.* 2014 | Clades in a multi-locus phylogeny | 8 |
| Corcoran *et al.* 2014 | Clustering of haplotypes | 11 |
| Boykin *et al.* 2014 | Existing taxonomic distinction and clades in consensus tree | 39 |
| Fusinatto *et al.* 2013 | Clades in a multi-locus phylogeny | 6 |
| Pérez *et al.* 2013 | ITS clades | 2 |
| Parmakelis *et al.* 2013 | Clades in a multi-locus phylogeny | 16 |
| Doyle *et al.* 2013 | Clades supported in $\geq 3$ of 4 gene trees | 16 |
| Pino-Bodas *et al.* 2013 | Existing taxonomic distinction and morphological differences | 9 |
| Prévot *et al.* 2013 | Clades in COI gene tree or combined mitochondrial tree | 8 |
| Zhao *et al.* 2013 | Existing taxonomic distinction | 6 |
| Cesar S. Herrera 2013 | Existing taxonomic distinction and geographic distribution | 4 |
| Niemiller *et al.* 2013 | Existing taxonomic distinction and geographic distribution | 4 |
| Hendrixson *et al.* 2013 | Existing taxonomic distinction and clades in mitochondrial gene tree | 4 |
| Keith & Hedin 2012 | Existing taxonomic distinction and geographic distribution | 76 |
| Niemiller *et al.* 2012 | Species delimitation/assignment via Brownie | 19 |
| Ardila *et al.* 2012 | Existing taxonomic distinction | 13 |
| Walker *et al.* 2012 | Existing taxonomic distinction | 11 |
| Donald *et al.* 2012 | Existing taxonomic distinction and geographic distribution | 9 |
| Walstrom *et al.* 2012 | Geographic distribution | 7 |
| Schmidt-Lebuhn *et al.* 2012 | Existing taxonomic distinction | 6 |
| Escobar *et al.* 2012 | Existing taxonomic distinction | 6 |
| Medina *et al.* 2012 | Morphological differences | 4 |
| Salariato *et al.* 2012 | Existing taxonomic distinction | 2 |
| Pettengill & Moeller 2012 | Existing taxonomic distinction | 2 |
| Levsen *et al.* 2012 | Existing taxonomic distinction | 2 |
| Willyard *et al.* 2011 | Existing taxonomic distinction | 32 |
| Gazis *et al.* 2011 | Clades in consensus tree | 16 |
| Taole *et al.* 2011 | Clades in ITS gene tree | 14 |
| Sakalidis *et al.* 2011 | Existing taxonomic distinction and morphological differences | 8 |
| Weisrock *et al.* 2010 | Existing taxonomic distinction and geographic distribution | 16 |
| Faustová *et al.* 2010 | Existing taxonomic distinction and geographic distribution | 8 |
| Groeneveld *et al.* 2010 | Existing taxonomic distinction and morphological differences | 4 |
| Valcárcel & Vargas 2010 | Existing taxonomic distinction | 4 |
| Koopman & Baum 2010 | Existing taxonomic distinction | 3 |
| Costanzo & Taylor 2010 | Existing taxonomic distinction | 2 |

When compared to other widely used species delimitation methods (Table 2) the *gsi* has many desirable properties. As well as having power to detect recently diverged lineages, the *gsi* differs from many comparable methods in not needing, as input parameters, the values of often difficult-to-estimate quantities such as the effective population size or mutation rate of the genetic sequences under consideration. The *gsi* value obtained for a given group is also purported to be comparable to those obtained for different groups within the same tree and between those arising from different studies (Cummings *et al*. 2008).

**Table 2.** “*θ*” refers to the population mutation rate for each locus, *“τ”* to the over-all height of a species tree in coalescent units. For DISCUSS, the ω parameter specifies a prior on the on the number of species present in a dataset, and *λ* and *μ* represent the speciation and extinction rates of a birth-death processes generating the underlying species tree. The description of the popABC approach to species delimitation refers specifically to the method employed by Camargo et al (Camargo *et al*. 2012)

| Method | Primary input | Group assignment | Other parameters | Result |
| --- | --- | --- | --- | --- |
| DISSECT (Jones *et al.* 2015) | Alignments | Inferred | $\theta, \tau, \omega, \lambda, \mu$ | Posterior distribution of model-paramaters, including species-tree and species- delimitation |
| BPP (Yang & Rannala 2010, 2014) | Alignments | *A priori* | $\theta, \tau$, species tree* | Posterior probability that species tree nodes represent speciation events |
| popABC (Lopes *et al.* 2009; Camargo *et al.* 2012) | Alignments | *A priori* | $\theta, \tau$, species tree | Posterior probability that species tree nodes represent speciation events |
| SpedeSTEM (Ence & Carstens 2011) | Gene trees | *A priori* | $\theta$ | Likelihood for population-delimitation models |
| Brownie (O’Meara 2010) | Gene trees | Inferred | None | Joint inference of maximum likelihood species-tree and species-delimitation |
| GMYC (Pons *et al.* 2006) | Gene tree | Inferred | None | Likelihood for species-delimitation models |
| *gsi* [14] | Gene tree | *A priori* | None | Statistic and hypothesis-test measuring each group’s phylogenetic exclusivity |

\* Note the species tree is an optional input parameter for BPP

In applying the *gsi* to empirical data, however, we have found the value of this statistic to be highly dependent on the structure of the tree outside of the group of interest. This dependence is not reflected in the way the statistic is typically applied and interpreted in species-delimitation studies, and this mismatch between the *gsi* as it is used and goals of species delimitation undermines the many advantages of the statistic.

## *The* gsi *measures exclusivity relative to the entire tree*

An example serves to illustrate the dependence of the *gsi* obtained for a particular group on the over-all structure of the phylogeny from which it is calculated. Take the tree presented in Figure 1. To calculate *gsi* for the group “a” in this tree we first need to calculate the intermediate statistic gs, which is defined as

**Figure 1.**
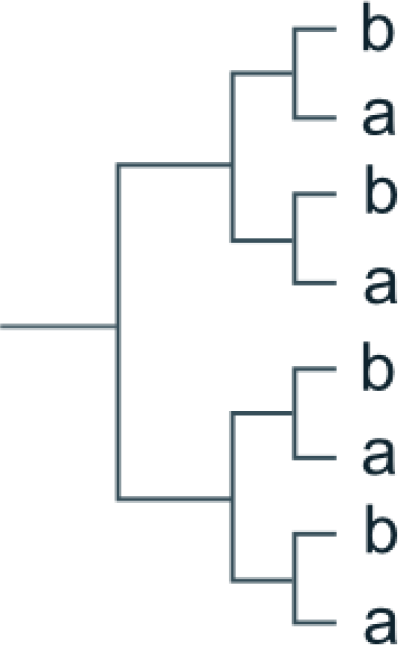
Hypothetical phylogenetic tree, with tips assigned to two groups “a” and “b”

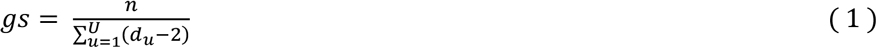

where *d_μ_*is the degree (i.e. the number of connections) of the node *u,* which is one of *U* total nodes in the smallest sub-tree uniting all members of a group and *n* is the minimum number of nodes that could be used to unite this group, which is one less than the number of leaves. In the case of Figure 1, *n* is 3, but all 7 nodes in the tree are needed to create a sub-tree uniting group “a”. As the tree is fully dichotomous, the degree of each node is 3. Thus, *gs* is 3 / (7 × (3-2)) = 3/7. To obtain *gsi* for group “a” the observed value of *gs* is normalised using the maximum and minimum obtainable value for the statistic given the size of the group and the number of nodes in the tree:

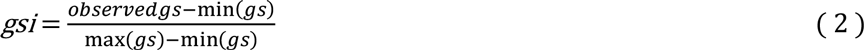

Here max(*gs*) is 1 (the case in which a group is united by the minimum number of nodes, i.e. monophyly) and min(*gs*) is given by the equation

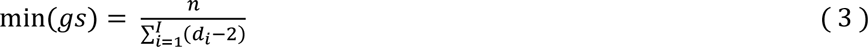

where *d_i_*is the degree of node *i*, one of *I* nodes in the entire tree. Because the smallest sub-tree uniting the group “a” in Figure 1 is the entire tree, min(gs) for this group is equal to the observed *gs* value. Thus, the numerator in equation (2) is 3/7 - 3/7 = 0 and so the value of *gsi* is also 0. This result is desirable, as the tree presented in Figure 1 has each group arranged in the least exclusive fashion possible. Nevertheless, defining min(gs) in this way means the value of *gsi* is partially dependent on the degree to which other groups in the tree fall into exclusive regions.

Consider now the tree presented in Figure 2, which could be obtained from genetic data underlying the topology illustrated in Figure 1 by simply adding further data from two distantly related taxa. Because the clade containing the “a” and “b” groups is unchanged the observed value of *gs* for “a” is still 7. The addition of the two groups “c” and “d” to the tree, however, has added 8 new dichotomous nodes. Thus the value of min(gs) is now 3/(15×(3-2)) = 3/15 and, following equation (2), *gsi* is equal to [3/7 - 3/15-3/15] ≈ 0.29. This difference arises from the inclusion of min(gs) in the calculation of gsi, which makes a *gsi* value obtained for a group a reflection of that group's exclusivity relative to the entire tree. This property is not desirable for a species-delimitation statistic, as it means large *gsi* values can be obtained for groups that do not represent a population that is divergent from all other samples in a given analysis. Moreover, it also compromises comparisons between groups within one study, or between values obtained from different studies.

**Figure 2.**
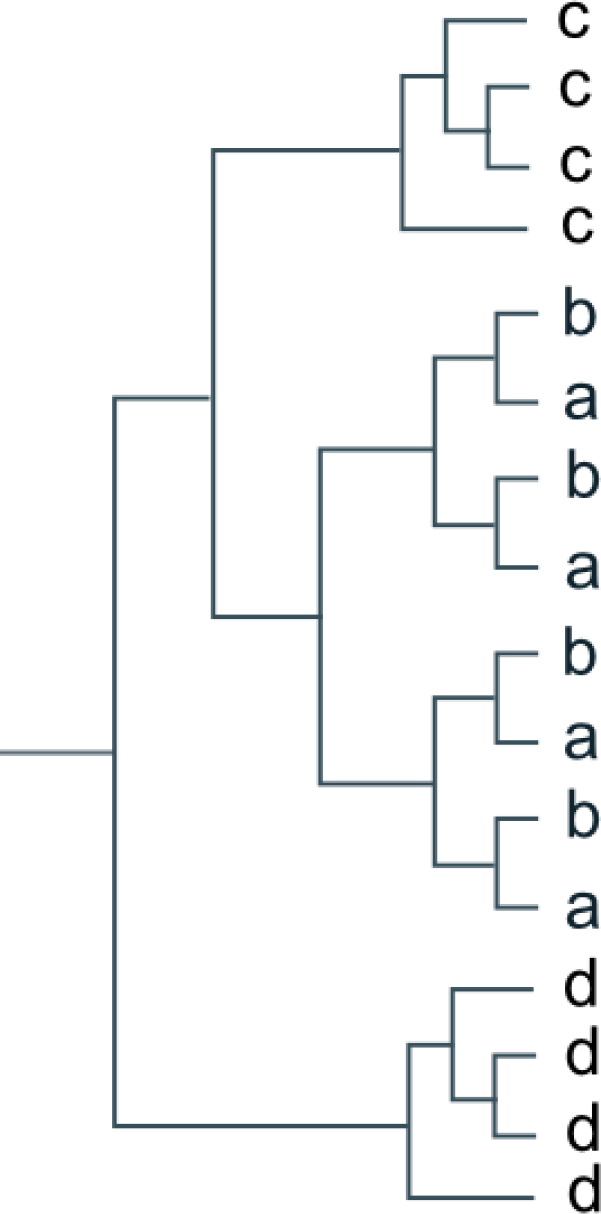
Hypothetical phylogenetic tree, obtained by adding two additional groups (“c” and “d”) to the tree presented in Figure 2.

A similar issue affects the hypothesis test that is often performed alongside the gsi. Statistical significance, or P-values, arising from this test are usually reported for each of several putative species under consideration in a single analysis. In practice, these P-values are interpreted as the results of independent tests that each group being considered is divergent from all other groups. In fact, as Cummings *et al*. (2008) make clear, because the test is performed by permuting group assignments across the entire tree, the null hypothesis being tested is that all individuals included in the tree come from a single panmictic population. It is seldom the case that all individuals considered in a species-delimitation study could plausibly have come from a randomly mating population. Thus, statistically significant results may simply represent the rejection of an implausible null hypothesis.

## *The* gsi *and species delimitation*

The problems discussed above are most likely to affect interpretation of the *gsi* when the statistic is calculated for a large number of groups, especially when those groups are likely to be divergent from at least some others under consideration. To determine how often the *gsi* has been used in such contexts, we performed a literature review (Table 1). We identified papers recorded as citing Cummings *et al*. (2008) in Web Of Science and Google Scholar. For each study we recorded the context in which the *gsi* was used, the largest number of groups considered in a single analysis and the criteria by which those groups were determined. The results of this analysis show the *gsi* has mainly been used in the context of species-delimitation (54 of 78 empirical studies) and that these studies have often applied the statistic to several groups (mean = 9.9, median = 6) for which there is *a priori* evidence for evolutionary divergence. The basis of the group assignment is frequently an existing taxonomic distinction, or a preliminary phylogenetic or clustering analysis performed on data from which the *gsi* was calculated. Worryingly, these circumstances are exactly those in which (as we show above) use of the *gsi* can be misleading. We did not find any papers in which the plausibility of the null hypothesis was considered in discussing the statistical significance of results.

To investigate how large the effect of including multiple divergent groups in the calculation of *gsi* is likely to be in practice, we calculated the statistic from simulated datasets. We used the program ms (Hudson 2002) to simulate gene trees arising from neutral evolution under the demographic history depicted in Figure 3, that is, four divergent populations with two (“a” and “b”) diverging at a time point *t* which was varied among simulations. We performed 500 simulations for each value of *t* between 0 and 1 *Ne* generations in 0.05 Ne increments, sampling 10 individuals per population in each simulation. For each simulation, we calculated the *gsi* for group “a” twice, first considering all populations in the simulation (the “four-population tree”) then after discarding individuals from the divergent populations “c” and “d” (the “two-population tree”).

**Figure 3.**
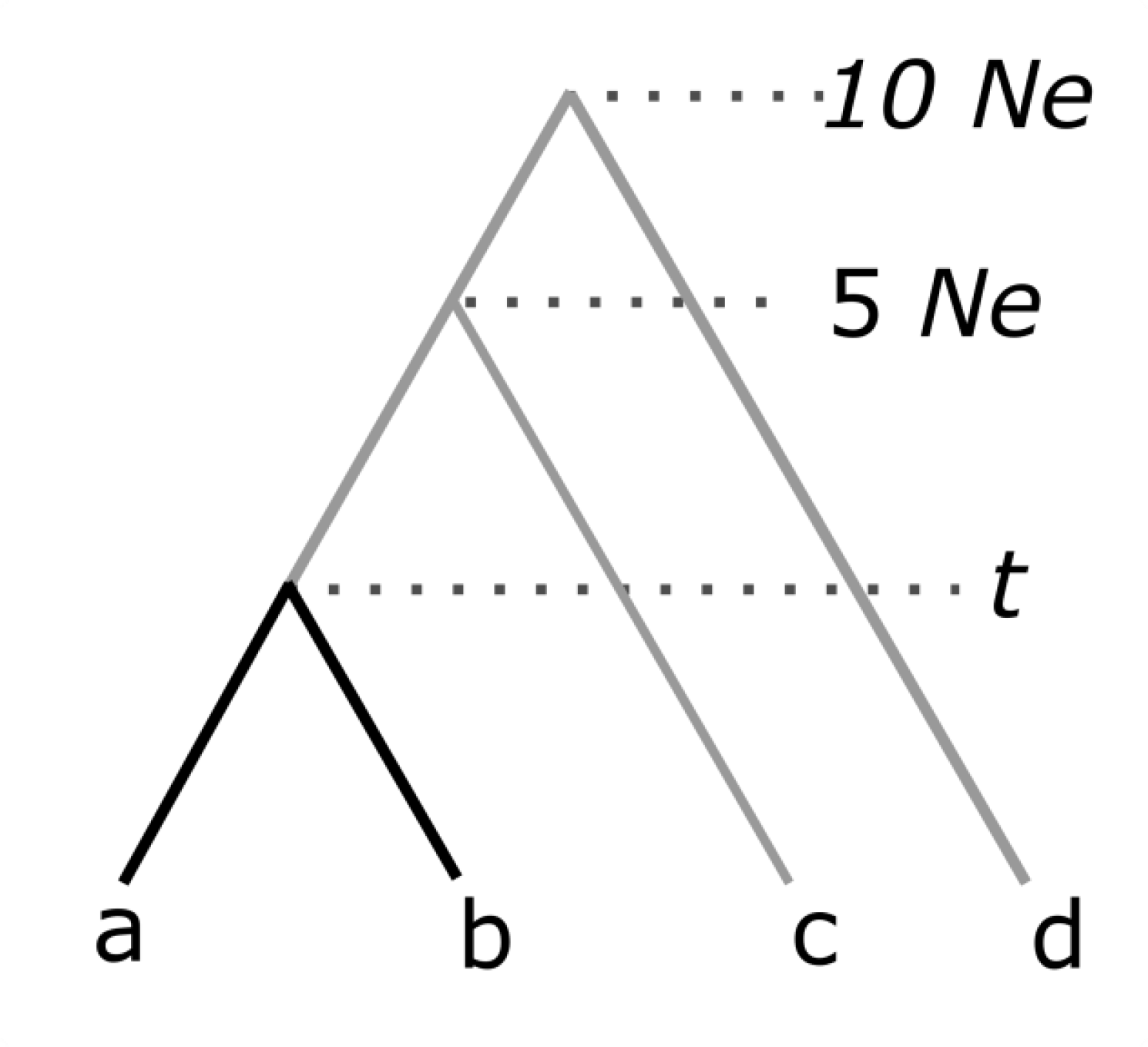
Demographic history under which simulations were performed. In each simulation *gsi* was calculating from a gene tree containing 10 individuals from each population (the four-population tree) and from a gene tree from which tips corresponding to individuals from populations “c” and “d” had been dropped (the two-population tree).

As expected, the mean *gsi* value obtained for the group “a” tracks the divergence of this population from population “b” in all simulations (Fig 4). However, the *gsi* value calculated from the four-population tree is substantially larger than the value obtained from the two-population tree, even though the same individuals make up the “a” population in each case. The difference is especially pronounced early in the divergence process, indeed, the expected value of *gsi* in the four-population case is high (0.40) even when *t* = 0 *Ne* (i.e. when populations “a” and “b” are panmictic with respect to each other). For every simulation, including those in which the “a” and “b” populations were panmictic, the calculation based on the four population-tree produced a significant result. By contrast, calculations based on the two-population tree produced a near-uniform distribution of P-values under panmixia and became increasingly likely to return significant results as the populations diverged.

**Figure 4.**
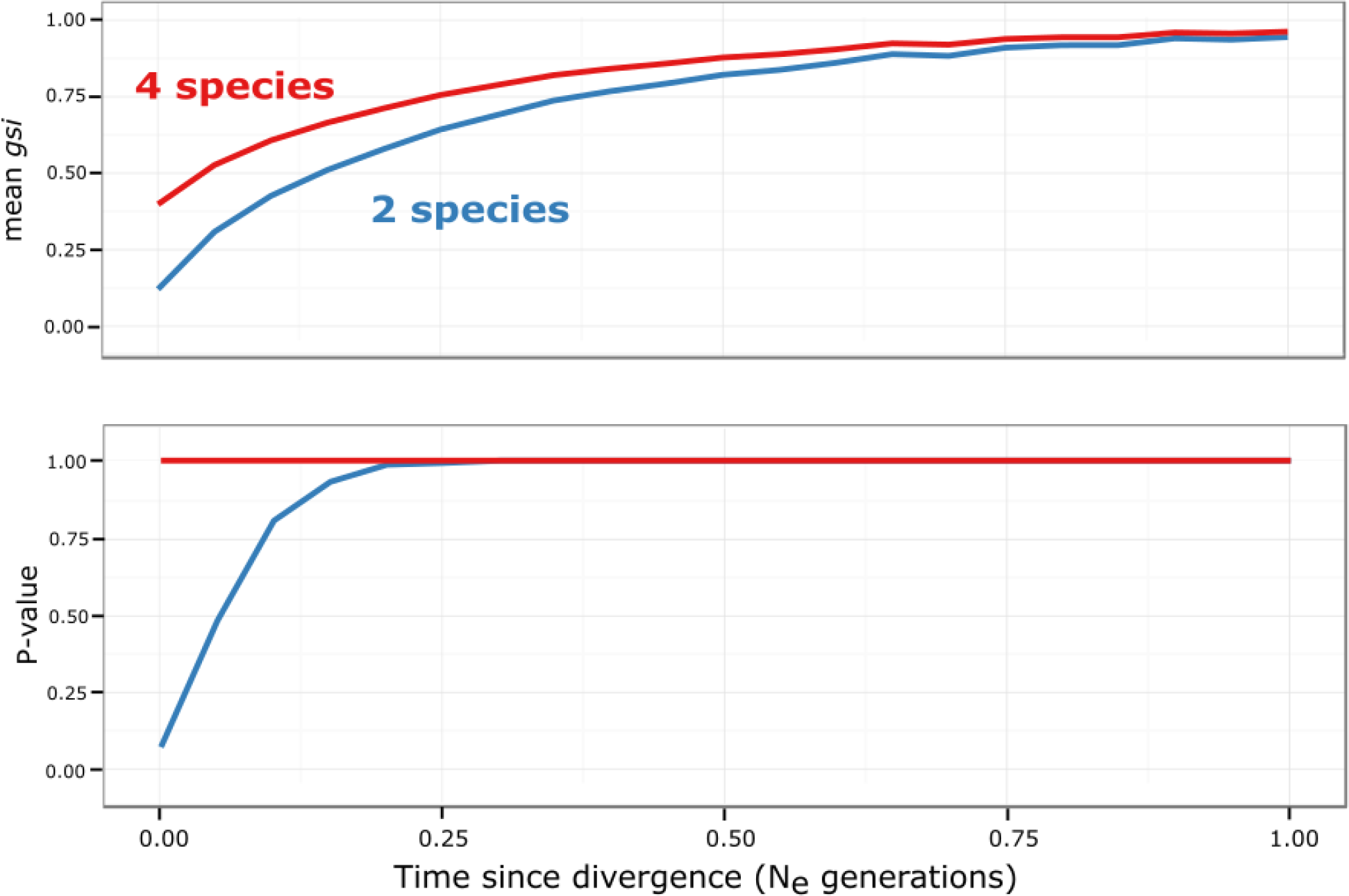
Above: The *gsi* value obtained for a group depends on the overall structure of the tree from which it is calculated. The red line represents the mean *gsi* value calculated for group “a” in the simulation depicted in Figure 3 when the all four populations are included in the calculation. The blue line is the mean *gsi* calculated for the same population when the divergent populations “c” and “d” are first discarded. Below: The hypothesis test usually performed alongside the calculation of *gsi* readily produces significant results for populations that are not divergent from all other populations. The red line represents the proportion of simulations in which population “a” was found to have a significant pattern of exclusivity in the four-population tree. The blue line represents the same proportion calculated from the two-population tree.

These simulations confirm that both the value and the significance obtained for the *gsi* of a given group is partially dependent on the degree to which other groups fall into exclusive regions of the phylogeny being considered. As we note above, this characteristic is not desirable in a statistic purported to relate only to the group under consideration, as it makes comparison of *gsi* values obtained from different trees problematic. Significant results can readily be obtained from populations that are not genealogically divergent from all others groups in an analysis.

## Aligning the gsi with species delimitation as it is practiced

As noted above, the *gsi* has many desirable properties (Table 2). Unlike many species delimitation methods, the *gsi* can be calculated without the need for often difficult-to-estimate population-genetic parameters as input. Additionally, the relative simplicity of the *gsi* means the statistic can be applied to large datasets. Unlike the GMYC (Pons et al., 2006), the *gsi* can be applied to unsorted gene trees and the *gsi* can be used to test the validity of proposed species suggested by morphological or other data. Given these unique properties of the *gsi,* we do no propose that empiricists discarding the statistic entirely. Rather, the properties described here should be carefully considered before the statistic before is applied to datasets.

There are likely to be multiple ways to reasonably incorporate the *gsi* in particular species delimitation studies; here we propose a general solution that retains the *gsi's* simplicity but removes its dependence on the overall structure of the tree from which it is calculated. Our proposed statistic, the “mean pairwise gsi” or *pwgsi* is calculated for a pair of groups, after all tips other than those in the groups of interest have been dropped from the phylogenetic tree under consideration. This approach can be applied to all putative species under consideration in a given study, or only to a subset that are of particular interest.

For example, to analyse the tree and group-assignments depicted in Figure 2 we first produce trees representing the possible pairwise comparisons of groups (Fig 5). For each tree, the *pwgsi* is simply the mean of the *gsi* values obtained for these two groups. Thus, in the case of Figure 5 the *pwgsi* for the “a:b” comparison is 0 and all other comparisons have *pwgsi* of 1. This approach requires at most 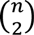values to be calculated, where *η* is the number of groups being considered. Thus, the pairwise approach is not subject to the computation limitations of methods that consider all possible partitions of a group-assignment (O’Meara 2010; Ence & Carstens 2011), and can be applied to datasets in which a relatively large number of groups are being considered. It also relatively easy to calculate the *pwgsi* with the existing genealogicalSorting software, as we demonstrate in Supplementary Text 1.

**Figure 5.**
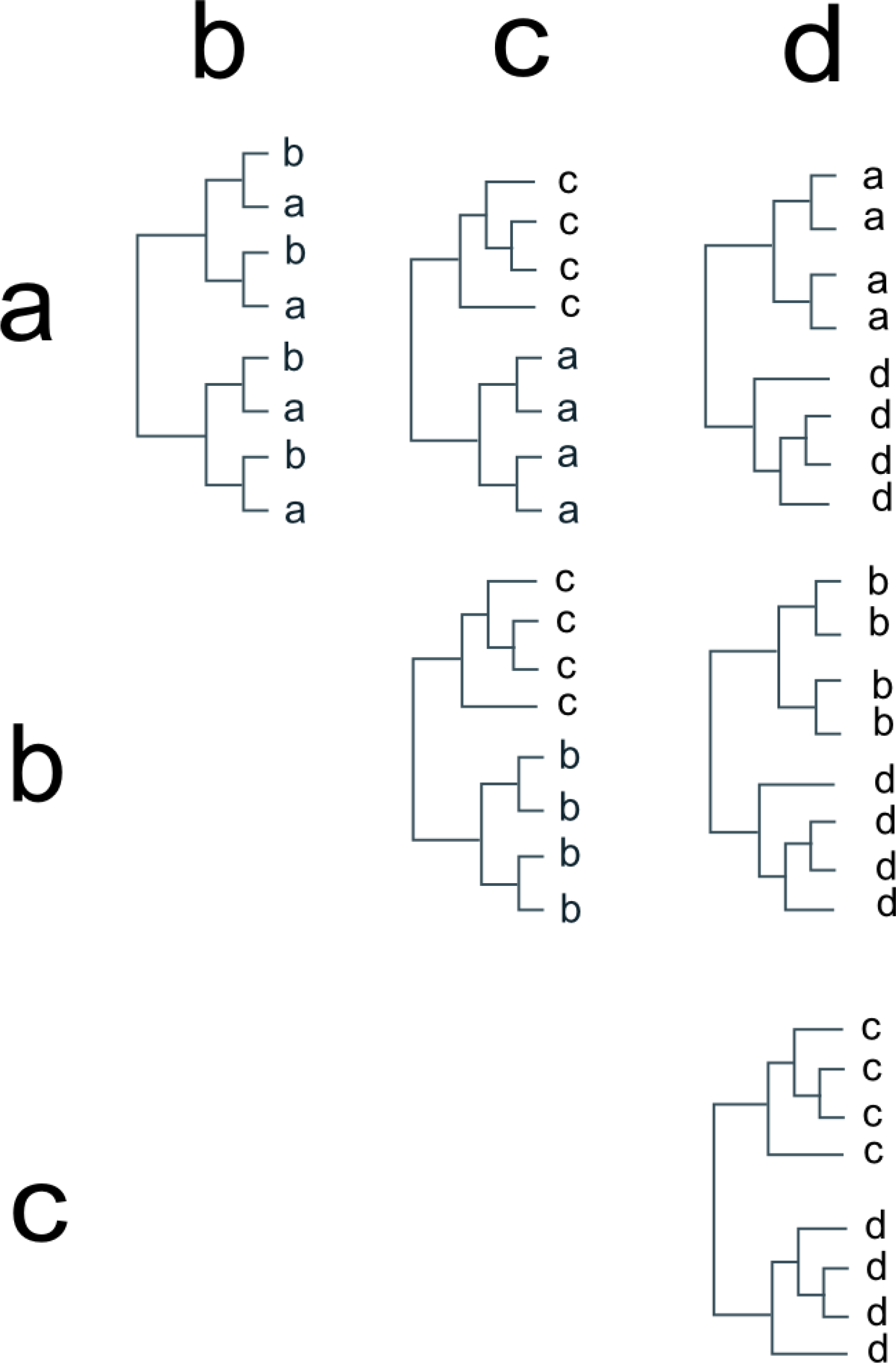
Trees used for pairwise comparison of all groups in Figure 2.

The pairwise approach aligns well with the goals of species delimitation studies, as the *pwgsi* quantifies each population's exclusivity relative to all other populations. Moreover, the pattern of *pwgsi* values resulting from a single analysis can identify groups that are not divergent with respect to each other, but are divergent from all other groups and thus might be considered part of a single divergent population in subsequent analyses (as is the case with “a” and “b” in example discussed above). This approach uses the same procedure as in the calculation of the two-population scores in Fig 4. We can infer from Fig 4, therefore, that the *pwgsi* tracks lineage divergence and a permutation test applied to a particular between-population comparison has a strong power to detect divergent groups.

## pwgsi *and population structure*

The high power of the *pwgsi* to detect an exclusive distribution of tips on a phylogeny may seem to make it an ideal statistic for species delimitation in the presence of incomplete lineage sorting. Care needs to be taken, however, in the interpretation of these results. The short period of divergence required to obtain significant results means that such results can be obtained even for what may turn out to be transient isolation between populations. Moreover, speciation is not the only one way in which a non-random distribution of groups might occur on a phylogeny. Specifically, sub-populations within a population with some degree of genetic structuring may be expected to fall into partially exclusive regions of a gene tree. To investigate the degree to which population structure affects the value of the *pw-gsi,* we simulated neutral gene trees under the scenario depicted in Figure 6. In this case, three populations diverged instantaneously at a time point that was held constant at 5 *Ne* generations. Two of the descendent populations (“a” and “b”) were united by ongoing and constant gene flow due to migration at a rate 4*N_e_m* with values {1, 2, 5, 25, 100}*. We performed 1500 simulations for each migration-rate value and sampled.

**Figure 6.**
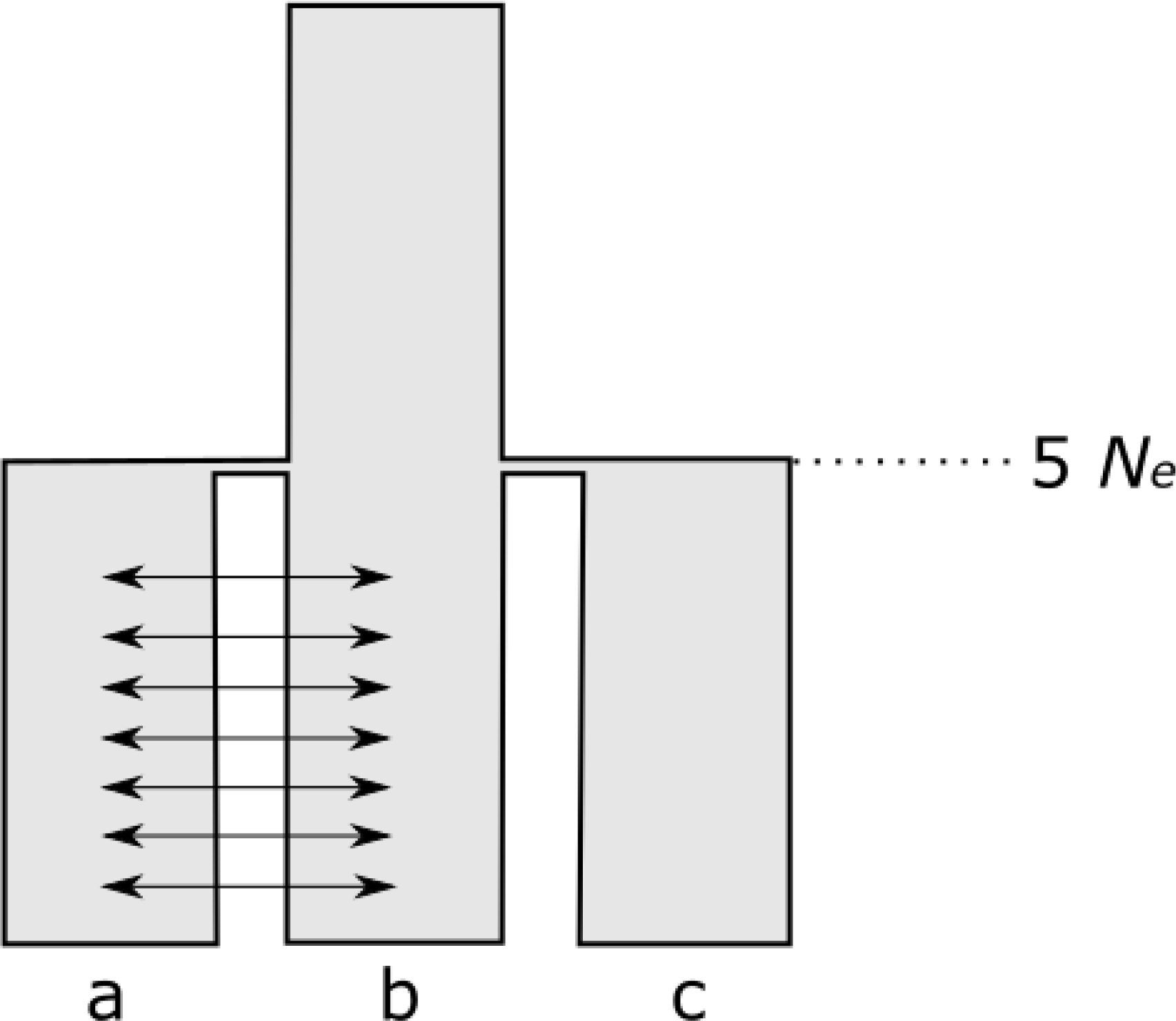
Demographic history under which population structure simulations were performed. Arrows represent ongoing gene flow between populations “a” and “b” after their divergence at 5 *Ne* generations.

The *pwgsi* value increased for the “a:b” comparison as the number of migrants exchanged between these populations decreased (and thus the populations became more structured) (Fig 7). We also investigated the power of the *pwgsi* to detect population structure in these simulations by performing 10^4^ group-label permutations per simulation. Significant results were obtained even with relatively limited population structure (Fig. 7). For example, when 4*N_e_m* = 100, a value giving a negligible expected *F_ST_* of < 0.01 (Wright 1949), more than 10% of simulations produced a result with a P-value < 0.05.

**Figure 7.**
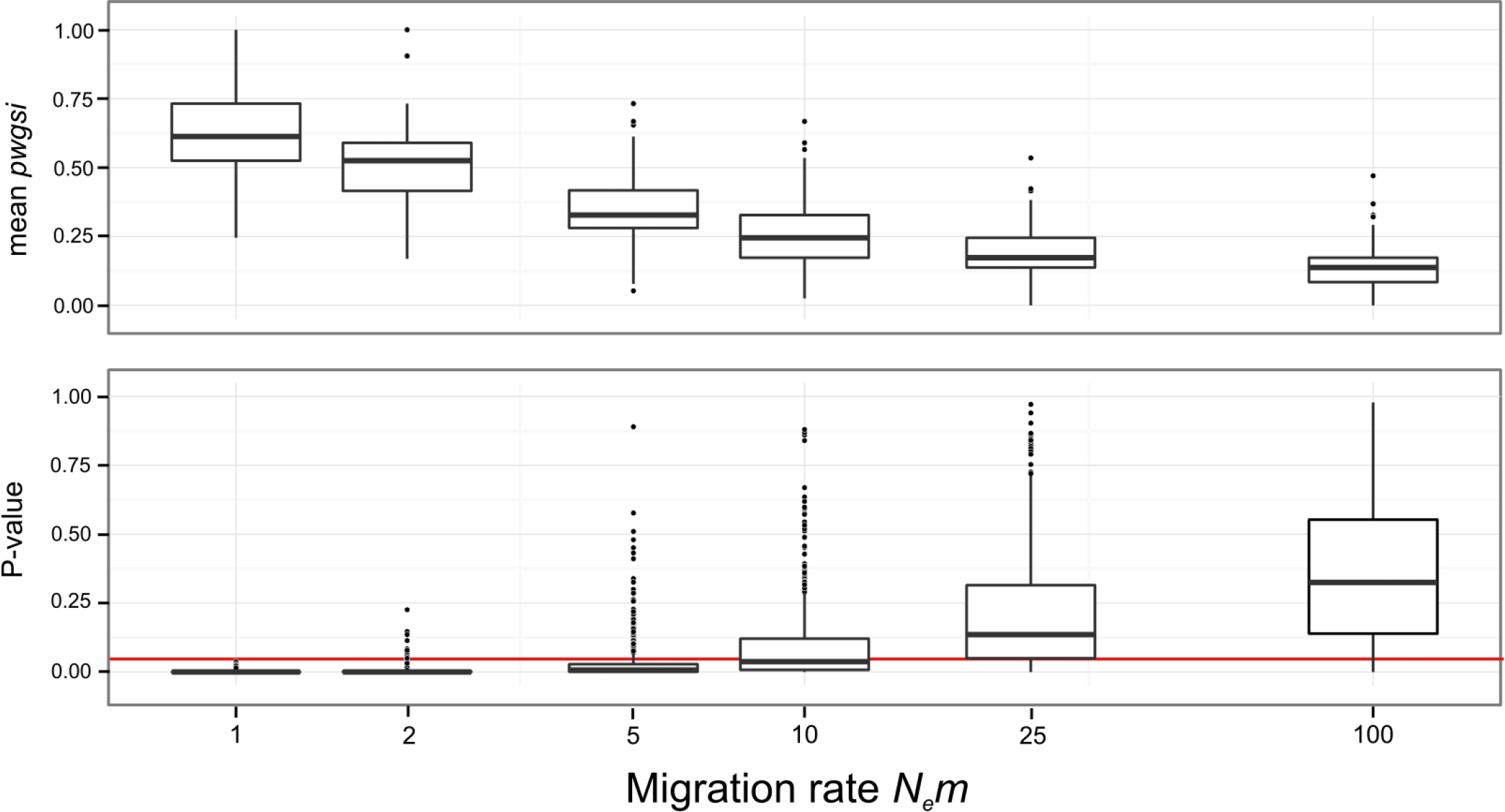
The *pwgsi* measures population structure as well as lineage divergence. Above: In the top panel, boxes represent distributions of *pwgsi* for the groups “a” and “b”, which were sampled from populations experiencing gene flow at a constant rate which varies along the x-axis (large values of *N_e_m* correspond to more migration and thus less population-structure). Below:. Boxes represent distributions of P-values obtained for the “a:b” comparison; red line the drawn at *P* = 0.05.Note, note x-axis is on a log-10 scale.

Although speciation with gene flow is certainly possible (Emelianov *et al*. 2004; Davison *et al*. 2005; Niemiller *et al*. 2008; de León *et al*. 2010) and perhaps even common (Nosil 2008), it is generally accepted even a small number of successful migrants are enough to prevent speciation in the absence of selection (Slatkin 1995; Gavrilets 2000). Speciation is only possible with greater rates of migration when very strong divergent selection is acting (Felsenstein 1981; Kirkpatrick & Ravigné 2002). Clearly then, the results of the *pwgsi* cannot be treated as unambiguous evidence that the groups being considered are different species. Rather, researchers need to consider it in the design of their studies and the interpretation of results. In particular, the *pwgsi* may be a poor choice of statistic if a putative species is known to have a distinct geographic distribution with respect to others to which it is being compared (or if the population samples being analysed come from different regions).

By contrast, our finding that the *pwgsi* measures population structure may make it a useful statistic for within-species phylogeographic studies, complementing the AMOVA approach (Excoffier *et al*. 1992) that is currently widely used for sequence data in this context. Indeed, the *gsi* has already been used in this context (Chen & Hare 2011; Gustafsson & Olsson 2012).

## Conclusions

Genetic sequences are a potentially powerful source of data for the discovery and delimitation of species. The results reported above, however, emphasise the care that needs to be taken in interpreting the results of DNA-based species-delimitation methods. We have shown that a naïve interpretation of gsi, a statistic that has been widely used in species-delimitation studies, can lead to erroneous conclusions. Although the *gsi* remains powerful approach to species when applied carefully (as in the *pwgsi* described here) it is still possible to obtain large *gsi* values and statistically significant results from populations connected by substantial gene flow.

Thus, we suggest that this statistic and other species-delimitation methods should be used as part of a genuinely integrative approach to taxonomy. In particular, the phylogenetic and population-genetic signals measured by species-delimitations methods should considered within the biological and ecological setting of a study, rather than as final arbiters of species’ status applied to hypotheses generated by other data.

## Acknowledgments

We thank three anonymous reviewers, Steffen Klaere, Giulio Dalla Riva, and Michael Cummings, whose comments on this manuscript greatly improved it. SK in particular should be credited with the insight described in the footnote on pgX.

## Footnotes

* Note, the inclusion of population “c” in this design illustrates the importance of the pw-gsi approach to quantifying lineage divergence. As this simulation proceeds population “c” is expected to become increasingly exclusive in gene trees arising from this history; thus the gsi values of “a” and “b” will increase over the course

